# Dietary protein from different sources escapes host digestion and is differentially modified by gut microbiota

**DOI:** 10.1101/2024.06.26.600830

**Authors:** Ayesha Awan, Alexandria Bartlett, J. Alfredo Blakeley-Ruiz, Tanner Richie, Casey M. Theriot, Manuel Kleiner

**Author notes:** Correspondence should be addressed to: Ayesha Awan - Manuel Kleiner.

## Abstract

Protein is an essential macronutrient and variations in its source and quantity have been shown to impact long-term health outcomes. Differential health impacts of dietary proteins from various sources are likely driven by differences in their digestibility by the host and subsequent availability to the intestinal microbiota. However, our current understanding regarding the fate of dietary proteins from different sources in the gut, specifically how component proteins within these sources interact with the host and the gut microbiota, is limited. To determine which dietary proteins are efficiently digested by the host, and which proteins escape host digestion and are used by the gut microbiota, we used high-resolution mass spectrometry to quantify proteins that constitute different dietary protein sources before and after digestion in germ-free and conventionally raised mice. We detected proteins from all sources in fecal samples of both germ-free and conventional mice suggesting that even protein sources with high digestive efficiency make it to the colon where they can serve as metabolic substrate for gut microbiota. Additionally, we found that specific component proteins of dietary protein sources were degraded to a greater extent in the presence of the microbiota. We found that specific proteins with functions that could potentially impact host health and physiology were differentially enriched in germ-free or conventionally raised mice. These findings reveal large differences in the fate of dietary protein from various sources in the gut which could explain some of their differential health impacts.

## Introduction

Dietary protein source has been shown to impact human health outcomes in large cohort studies. In these studies, dietary proteins of animal origin, such as those found in eggs and beef, were associated with increased all-cause mortality, and substituting them with plant-sourced dietary protein reduced mortality^1–4^. Additionally, increased animal protein intake also enhances the risk and incidence of inflammatory bowel disease (IBD) and contributes to relapse in ulcerative colitis^5,6^. While the exact mechanisms by which dietary protein from various sources impacts the host are not well understood, interactions between undigested dietary protein and the intestinal microbiota are thought to play a key role^7,8^. Specific dietary components, including dietary protein and fiber, have been shown to impact the composition and function of the gut microbiota^9–11^. Dietary proteins that escape host digestion can be converted by the gut microbiota to produce metabolites that impact host health, such as beneficial short-chain fatty acids or proinflammatory ammonia, amines, and hydrogen sulfide^7,12^. The digestibility of proteins is an important determinant of how much and which dietary proteins escape host digestion and are accessible to the gut microbiota in the large intestine. Animal and plant-sourced proteins have different digestibility and some animal proteins are considered to be more bioavailable and easier to digest^13,14^. Hence more plant protein is expected to escape host digestion and be metabolized by the gut microbiota. However, which specific protein components of dietary proteins from various plant and animal sources escape host digestion and undergo microbial degradation in the large intestine remains unknown.

Current methods that investigate dietary protein digestibility are limited in their ability to address questions regarding the fate of individual dietary proteins as they pass through the intestinal tract because they rely on amino acid, nitrogen, or crude protein quantification^15–18^. These methods are unable to distinguish between host, microbial, and diet-sourced proteins and cannot provide details regarding the interactions of individual proteins in dietary protein with the host and the gut microbiota. Furthermore, amino acid, nitrogen, and crude protein quantification do not account for the impacts of proteins that retain function in the intestinal environment, such as anti-nutritional factors like trypsin inhibitors^16,17,19^. Shotgun proteomics is an alternative method that can be used to address these knowledge gaps because it quantifies the individual dietary proteins in diet and fecal samples and is able to distinguish dietary, host and microbial proteins in these samples^20^.

Here, we used proteomics to investigate 1) how dietary proteins from various plant and animal sources vary in their composition, 2) which individual proteins from a specific protein source are efficiently digested by the host, and 3) which proteins escape host digestion and are accessible to microbial degradation in the large intestine. We determined the composition and fate of dietary proteins from six different plant and animal sources, including soy, casein, brown rice, yeast, pea, and egg white in the presence and absence of the gut microbiota. These dietary protein sources were selected based on their relevance to human and animal nutrition. Casein is the primary protein in dairy milk and egg white is considered a rich source of high quality protein^13,14^. Soy is a complete protein, containing all essential amino acids and is widely used in plant based vegetarian and vegan diets^21,22^. Pea and brown rice protein are popular in plant-based protein diets and supplements^23,24^. Casein and soy are also the primary protein sources in the standard chow and defined diets usually used in mouse studies^25^. Our results show that dietary proteins from each of these six sources vary in terms of their composition and complexity, as well as their digestive efficiency in the host and accessibility to and use by the gut microbiota.

## Methods

### Animals and housing

Animals were obtained and housed as described in Bartlett et al.^28^. For this study, we used 12 germ-free (6 male, 6 female, NCSU gnotobiotic core) and 12 conventionally raised (6 male, 6 female, Jackson Labs, Bar Harbor) C57BL/J6 mice, aged 3-6 months. The germ-free mice were kept in gnotobiotic isolators throughout the experiment. They were regularly monitored for sterility using anaerobic culturing techniques. The male and female mice were housed separately in groups of three with a 12-hour dark/light cycle at an average temperature of 70°C and 35% humidity. The bedding was autoclaved and sterile water (Gibco) was used for both germ-free and conventional mice. The diets for both germ-free and conventional mice were gamma-irradiated and for the germ-free mice, the outside of the vacuum sealed diet packages was further sterilized using a disinfectant before introduction to the cages. Cage changes for the conventional mice were performed in a laminar flow hood. The animals were fed, weighed, and assessed daily by trained animal handlers. All animal experiments were conducted in the Laboratory Animal Facilities at the NCSU College of Veterinary Medicine, which is accredited by the Association for the Assessment and Accreditation of Laboratory Animal Care International (AAALAC). The animal care and use protocol was approved by NC State’s Institutional Animal Care and Use Committee (Protocol # 18-034-B for conventionally raised mice and # 18-165-B for germ-free mice).

### Animal diets and sample collection

The diets used in this study were defined diets obtained from Inotiv (previously Envigo Teklad). The composition of the diets in terms of protein, carbohydrate, fat, vitamin and mineral components was controlled across the different diet groups (Supplemental Table 4). Components other than the source of the dietary protein did not change and were consistent across the different diets. Diets were gamma-irradiated and vacuum packaged so that they could be sterilized on the outside while introducing them into the gnotobiotic isolators. The diets contained purified protein from a single source and were not supplemented with amino acids. We fed the mice a sequence of diets containing 20% soy, 20% casein, 20% brown rice, 40% soy, 20% yeast, 40% casein, 20% pea, and 20% egg white. Diets were available to mice *ad libitum* for 7 days per diet (Fig. 1A). At the end of the dietary sequence mice were divided into two groups where one group was fed 20% casein and the other was fed 20% soy again as a control. Fecal samples were collected from each mouse on the seventh day after feeding the defined diet for a week before switching to the next diet. Different diets were fed to the same mice to minimize the number of mice needed for ethical reasons and to account for interindividual microbial variability, which can be as high as 45%^48^. Additionally, based on previous studies that have shown that the gut microbiota changes robustly and reproducible in response to dietary changes in as few as 3 days, the order of diets was not considered to have any lingering or compounding effects 7 days after diets were switched^10,49^. We confirmed this by re-feeding the soy and casein diets at the end of the feeding regimen as controls and found that the gut microbiota returned to the same state observed in weeks 1 and 2 when soy and casein were initially fed^11^. We initially also had chicken bone broth as a third animal protein source in the feeding regimen. However, mice showed signs of clinical sickness on that diet, including weight loss by the third day, so we stopped feeding it and fed standard chow for the remainder of the seven days. Since no data was collected for the chicken bone broth diet, it is not included in the analysis. At the end of the dietary rotation the mice were humanely euthanized by CO_2_ asphyxiation and intestinal contents from the duodenum, ileum, cecum and colon of each mouse were collected. Fecal and intestinal content samples were placed in NAP preservation solution (935 ml autoclaved MilliQ water, 700 g Ammonium Sulfate, 25 ml of 1 M Sodium Citrate and 40 ml of 0.5 M EDTA adjusted to pH 5.2 using 1 M H_2_SO_4_)^50^ at a 1:10, sample-to-solution ratio. This was done to prevent sample modification, for example by proteases, during the prolonged time that it takes to remove samples from the gnotobiotic isolators. For consistency, samples from germ-free mice and conventionally raised mice were treated identically. Samples were roughly homogenized using a sterilized disposable pestle before being frozen at −80°C. All mice were handled following the approved IACUC protocol.

**Figure 1.**
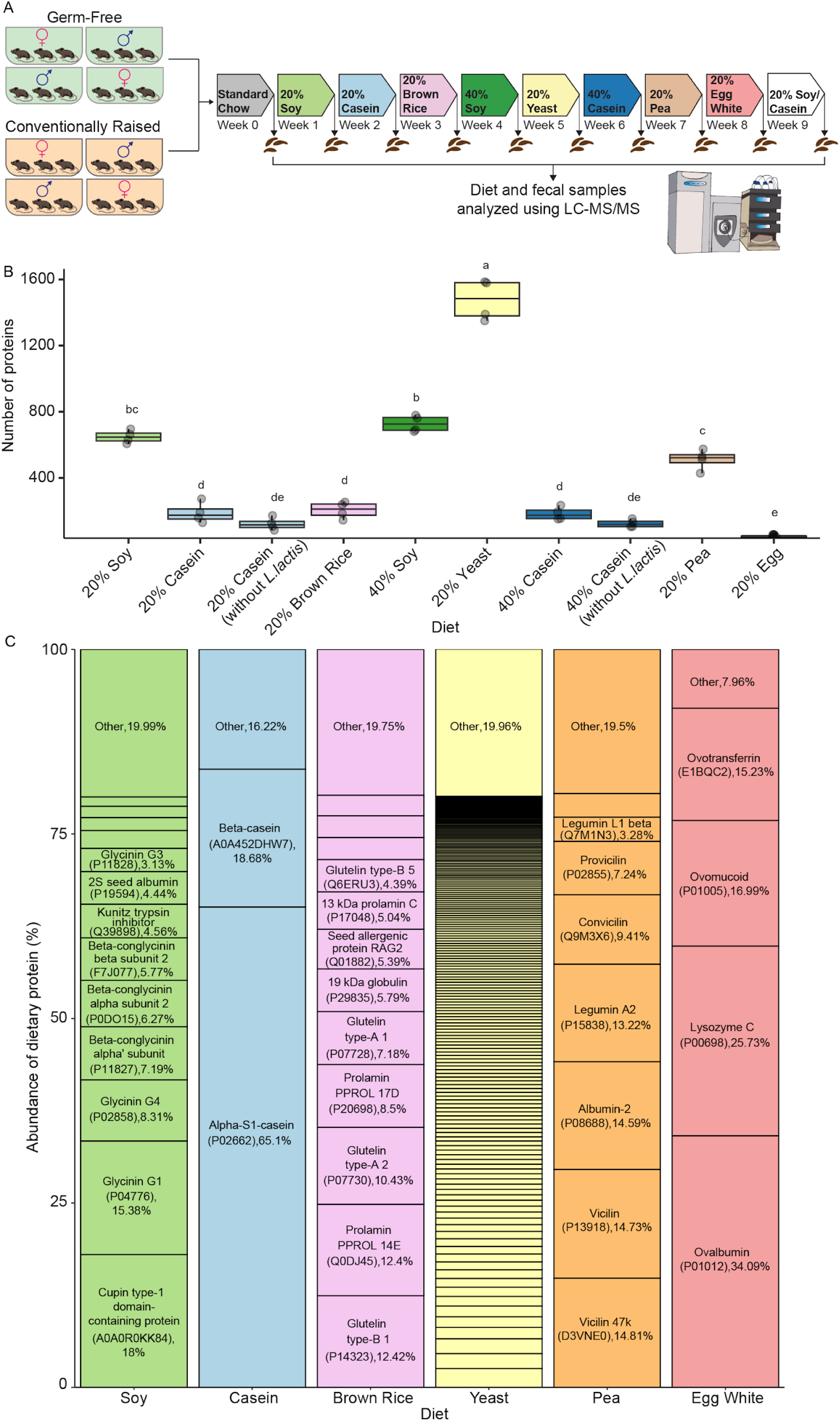
Sources of dietary protein differ in composition and complexity. A) Schematic showing the experimental design. Each mouse was fed the defined diet containing purified dietary proteins from one of six sources for a week and fecal samples were collected at the end of each week. The diet and fecal samples were analyzed using LC-MS/MS. Colors depicting each of the dietary groups and the germ-free and conventionally raised mice are used consistently throughout the manuscript. B) Boxplot showing a comparison of the number of individual proteins detected in each dietary protein source (4 replicates per source) using proteomics. The letters above the boxplots indicate statistical differences, where boxes with the same letter are not significantly different (ANOVA followed by a Tukey’s HSD pairwise comparison, *p*-value < 0.05). C) Plot showing the relative abundance of component proteins in each of the dietary protein sources (ranging from the most abundant proteins at the bottom to the least abundant proteins at the top). Labels consist of the name of the protein, the protein accession from Uniprot, and the average relative abundance of the protein across 4 replicates. Names and accession numbers of the unlabeled component proteins can be found in Supplemental Table 1.

### Amino Acid Composition Analysis

Amino acid analysis on all diets except for brown rice was conducted by Eurofins Scientific Inc., Des Moines, USA (Accreditation: ISO/IEC 17025:2017) using the reference AOAC methods (AOAC International, 2012) 988.15 for tryptophan, 994.12 for cystine and methionine and 982.30 for other amino acids. Amino acid composition of the purified brown rice protein was obtained from a previous study that analyzed the exact same product (Oryzatein Silk 80™) used in our study^51^.

### Protein extraction from diet and fecal samples

Protein extraction and peptide preparation were conducted as described in Bartlett et al.^28^ We prepared peptides from food pellets and the collected fecal samples. For the fecal samples, we removed the NAP solution by centrifugation (21,000 x g, 5 min). We placed the food and fecal samples in Lysing Matrix E tubes (MP Biomedicals). For the food samples we used 100 mg (4 replicates from each source). SDT lysis buffer (4% (w/v) SDS, 100 mM Tris-HCl pH 7.6, 0.1 M DTT) was added to the samples and the samples were bead beat (5 cycles of 45 s at 6.45 m/s, 1 min between cycles). Samples were heated to 95°C for 10 minutes and the lysates were centrifuged (21,000 x g, 5 min). We followed the filter aided sample preparation (FASP)^52^ protocol for peptide preparation. We mixed 60 µl of the lysate with 400 µL of UA solution (8 M urea in 0.1 M Tris/HCl pH 8.5) and loaded the mixture onto a 10 kDa 500 µL filter unit (VWR International) and centrifuged at 14,000 x g for 30 minutes.

This step was repeated 3 times to load the filter to capacity. The filters were washed using 200 µl of UA solution and centrifuged at 14,000 g for 40 min followed by incubation with 100 µl IAA (0.05 M iodoacetamide in UA solution) for 20 min and centrifugation at 14,000 *g* for 20 min. Filters were then washed 3 times with 100 uL of UA buffer and 3 times using 100 µl of ABC (50 mM Ammonium Bicarbonate) buffer. The proteins were digested into peptides by adding 0.95 µg of MS grade trypsin (Thermo Scientific Pierce, Rockford, IL, USA) solubilized in 40 µl of ABC buffer to the filters and incubating for 16 hours in a wet chamber at 37 °C. Peptides were eluted by centrifugation at 14,000 x g for 20 minutes followed by another elution using 50 uL of 0.5 M NaCl and centrifugation at 14,000 x g for 20 minutes. Peptide concentrations were determined using the Pierce Micro BCA assay (Thermo Scientific Pierce) according to the manufacturer’s instructions.

### LC-MS/MS analysis of peptides from diet and fecal samples

Peptides from diet and fecal samples were analyzed using 1D-LC-MS/MS as previously described in Bartlett et al.^28^ The peptide samples were block randomized, and 600 ng for mouse fecal samples and 300 ng for diet samples were loaded onto a 5 mm, 300 μm ID C18 Acclaim® PepMap100 pre-column (Thermo Fisher Scientific) with loading solvent A (2% acetonitrile, 0.05% TFA) in an UltiMate™ 3000 RSLCnano Liquid Chromatograph (Thermo Fisher Scientific). An EASY-Spray analytical column heated to 60°C (PepMap RSLC C18, 2 µm material, 75 cm× 75 µm, Thermo Fisher Scientific) was then used to separate the peptides. A 140 min gradient at a flow rate of 300 nl/min was used for peptide separation where the first 102 minutes of the gradient went from 95% eluent A (0.1% formic acid) to 31% eluent B (0.1% formic acid, 80% acetonitrile), then 18 min from 31 to 50% B, and 20 min at 99% B. A 100% acetonitrile wash run was performed between each sample to minimize carryover. The eluted peptides were ionized using an Easy-Spray source and analyzed in a Q Exactive HF hybrid quadrupole-Orbitrap mass spectrometer (Thermo Fisher Scientific) with the following parameters: m/z 445.12003 lock mass, normalized collision energy equal to 24, 25 s dynamic exclusion, and exclusion of ions of +1 charge state. Full MS scans were acquired for 380 to 1600 m/z at a resolution of 60,000 and a max IT time of 200 ms and data-dependent MS^2^ was performed for the 15 most abundant ions at a resolution of 15,000 and max IT of 100 ms.

### Database construction and protein identification and quantification

A protein sequence database containing dietary, host, and microbial proteins was constructed as previously described in Bartlett et al.^28^ For the dietary components in the database, reference protein sequences for soy (*Glycine max*, UP000008827), casein (*Bos taurus*, UP000009136), brown rice (*Oryza sativa*, UP000059680), yeast (*Cyberlindnera jadinii*, UP000094389), pea (*Cajanus cajan*, UP000075243) and egg white (*Gallus gallus*, UP000000539) were downloaded from UniProt and added to the host and microbial components to construct separate databases for each diet. The diet-specific databases^53^ were used to search the MS^2^ spectra using run calibration and the Sequest HT node in Proteome Discoverer software version 2.3 (Thermo Fisher Scientific) with the following settings: trypsin (Full), maximum 2 missed cleavages, 10 ppm precursor mass tolerance, 0.1 Da fragment mass tolerance and maximum 3 equal dynamic modifications per peptide.

Dynamic modifications including oxidation on M (+15.995 Da), deamidation on N, Q, R (0.984 Da), and acetyl on the protein N terminus (+42.011 Da), and the static modification carbamidomethyl on C (+57.021 Da) were considered. The Minora Feature Detector node was used for protein quantification based on area under the curve, with the following parameters: 5 minimum number of non-zero points in a chromatographic trace, 0.2 min maximum retention time of isotope pattern multiplets, and high PSM confidence. The percolator node in Proteome Discoverer was used to calculate peptide false discovery rate (FDR) with the following parameters: maximum Delta Cn 0.05, a strict target FDR of 0.01, a relaxed target FDR of 0.05, and validation based on q-value. The precursor ion quantifier node was used for quantification using the following settings: unique and razor peptides used for quantification, and precursor abundance based on area. Only master proteins were included in the downstream data analysis.

### Data processing, statistical analysis, and data visualization

Total sum scaling was used to normalize the relative abundances of proteins. To determine the relative abundance (%) of dietary, host, and microbial proteins in the fecal samples, we summed the abundance of proteins within those categories. We used ANOVA followed by Tukey’s HSD post hoc comparison to determine differences in the relative abundance of dietary proteins between diet, germ-free mice and conventionally raised mice. To make the data comparable across dietary and treatment groups, and to address variations in total protein content, we normalized dietary protein abundances separately from host and microbial proteins using total sum scaling for further analysis. Dietary proteins present in at least 75% of samples in at least one group (group determined using mouse type and dietary protein source) were included in the downstream analysis. For the principal coordinate analysis and the PERMANOVA, we transformed the dietary protein data using a centered log-ratio transform using the compositions package (version 2.0.5)^54^ in R^55^. To identify individual proteins that were statistically different in abundance across diet and fecal samples we used the two-sample t-test with unequal variances (Welch’s t-test). To determine log fold-change differences in the abundance of individual proteins in the diet and fecal samples, we log_2_ transformed the normalized relative abundance values. To prevent infinite ratios during analysis, zeros in the data were replaced with a value equal to one-tenth of the smallest non-zero value in the dataset prior to calculating log fold-change. For visualization purposes, the proteins that were at least 1% abundant in either of the compared groups or had at least a 1 log-fold difference were selected. All p-values were corrected for multiple hypothesis testing using the Benjamini-Hochberg correction in the rstatix package^56^ in R. R (R version 4.2.2) and Excel were used for all data processing and analysis. R (ggplot2, version 3.5.1)^57^, Origin (version 2022b, OriginLab Corporation, Northampton, MA, USA) and Adobe Illustrator were used for data visualization.

## Results

### Purified dietary proteins comprise hundreds to thousands of proteins

We fed defined diets containing purified dietary protein from one of these six sources to male and female germ-free and conventionally raised mice in a weekly feeding regimen where the mice received a different diet each week. In total, we used eight different diets, which included diets with 20% protein content for all six dietary protein sources, and for casein and soy, we also used diets with 40% protein content. At the end of each week, we collected fecal samples from each mouse. We then analyzed the diet and fecal samples using liquid chromatography coupled high-resolution mass spectrometry and quantified hundreds to thousands of dietary proteins in each sample (Fig. 1A).

We used proteomics to analyze the protein content of each purified dietary protein source (Fig. 1A). Each dietary protein source was composed of tens to thousands of constituent proteins ranging from 44 proteins in the egg white protein diet to 1476 proteins in the yeast protein diet (Fig. 1B, Supplementary Data Table 1). Dietary proteins from animal sources (casein and egg white) had fewer individual proteins, and, as such, were less diverse than dietary proteins from plant sources (soy, pea, and brown rice). Yeast, defined as a dietary protein of microbial origin^26,27^, had the greatest number of constituent proteins. In some sources, such as casein and egg white, very few proteins made up the bulk (80%) of the dietary protein, while other dietary proteins, such as yeast, consisted of many low abundance proteins (Fig. 1C). In order to further determine differences and similarities among these protein sources, we analyzed their amino acid composition. The amino acid abundances for all protein sources were similar, deviating by no more than 30% from the mean abundance across all sources, with the exception of cystine, methionine, proline and tyrosine (Supplementary Figure 1, Supplemental Table S5). Egg white contained the highest amount of cystine and casein contained the highest amount of proline. Methionine was higher in casein and egg white compared to the other plant and microbial dietary protein sources. Brown rice contained the highest amount of tyrosine.

Casein protein consisted of bovine proteins, including alpha and beta-casein which constituted 80% of total protein abundance, as well as a large number of microbial proteins, specifically *Lactococcus lactis* proteins, which were likely remnants of the casein protein processing^28,29^. These *L. lactis* proteins (66 in the 20% casein diet and 59 in the 40% casein diet) were of low abundance and made up only 1% of the total protein abundance in the purified casein diet. Eighty percent of the egg white protein was comprised of ovalbumin, ovomucoid, lysozyme, and ovotransferrin. From plant sources, 8, 12 and 13 proteins made up 80% of the total protein in pea, brown rice, and soy protein diets, respectively. The seed storage proteins Vicillin (14.8%), Glutellin (12.4%), and Glycinin (18%) were the most abundant dietary proteins in the pea, brown rice, and soy protein diets, respectively. In addition to seed storage proteins, we also detected proteins previously shown to impact health, such as allergens like the seed allergenic protein (Q01882) in the brown rice protein diet^30^, and anti-nutritional factors like the Kunitiz trypsin inhibitor (Q39898) and lectin protein (P05046) in the soy protein diet^22,31^. In summary, these purified sources of dietary protein covered a large range in diversity based on the number of constituent proteins and their abundances.

### Different dietary protein sources are digested at varying efficiencies in the presence and absence of the gut microbiota

To determine the impact of the host and the gut microbiota on the digestion of different sources and amounts of dietary protein, we compared the abundance of dietary protein recovered in the fecal samples of germ-free and conventionally raised mice across the eight diets. Comparison across all eight diets showed that the fecal samples of germ-free mice fed the 20% brown rice protein diet had significantly higher dietary protein than the other diets (Fig. 2A). After the brown rice group, the 20% egg white and 40% casein groups had the highest amount of dietary protein in the feces, followed by the 20% casein group. The lowest amount of dietary protein was recovered in the fecal samples of germ-free mice fed the 20% soy, 40% soy, 20% yeast, and 20% pea protein diets.

**Figure 2.**
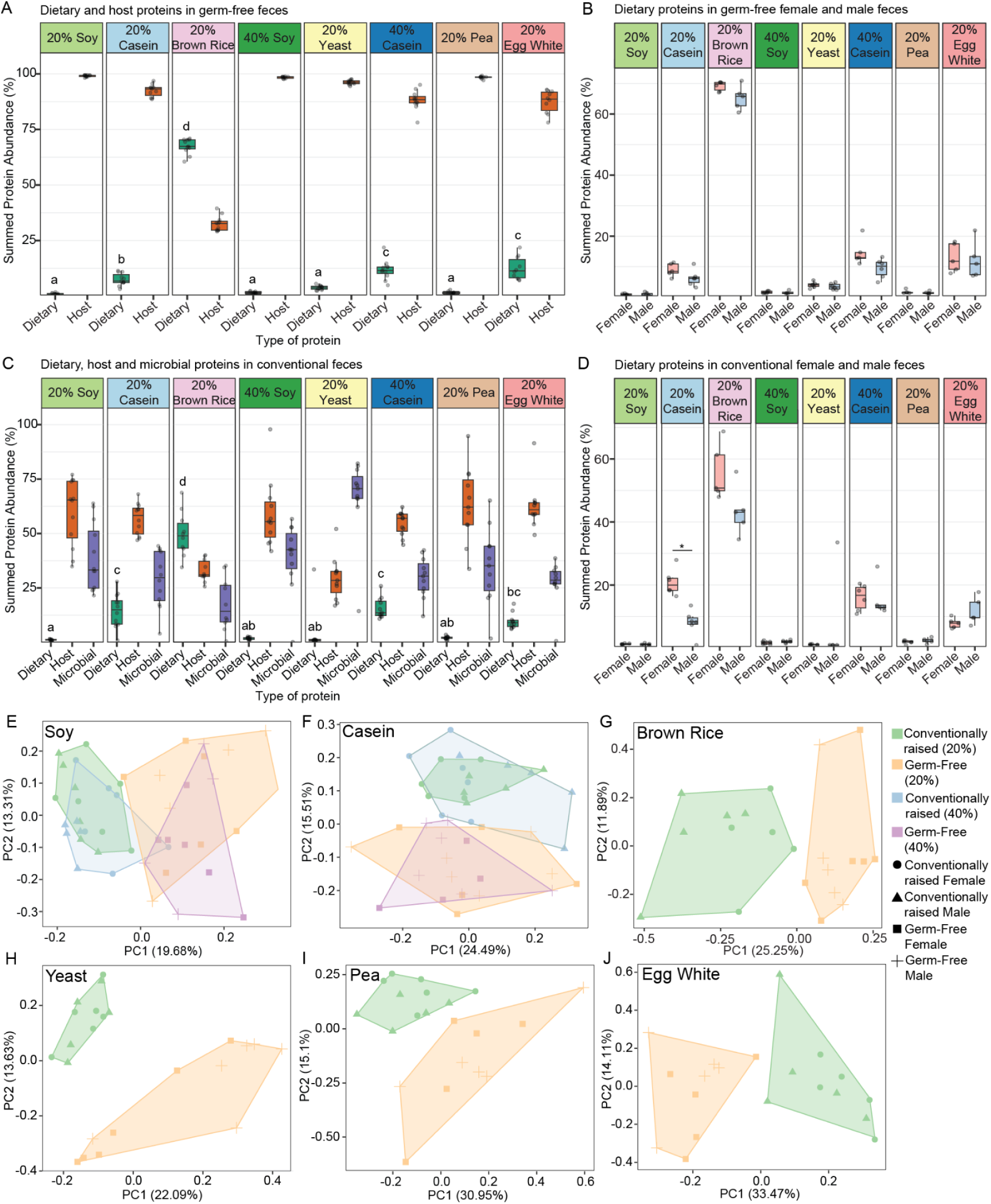
**Differential digestion of dietary protein sources in the presence and absence of a gut microbiota**. A) Boxplots showing the summed protein abundance in percentages of dietary (green) and host (orange) proteins in fecal samples of germ-free mice (20% soy (n=12), 20% casein (n=11), 20% brown rice (n=11), 40% soy (n=10), 20% yeast (n=12), 40% casein (n=10), 20% pea (n=10), and 20% egg white (n=10). The letters on top of the dietary (green) boxplots denote statistical differences between groups as determined by an ANOVA test followed by Tukey’s HSD post hoc comparison (p<0.05). Groups that have at least one similar letter are not significantly different from each other, while groups with different letters are significantly different. B) Boxplots showing abundance of dietary proteins in fecal samples of germ-free male (blue) and female (pink) mice. C) Boxplots showing the summed protein abundance in percentages of dietary (green), host (orange) and microbial (purple) proteins in fecal samples of conventionally raised mice (20% soy (n=12), casein (n=11), brown rice (n=10), 40% soy (n=11), yeast (n=11), 40% casein (n=11), pea (n=11), and egg white (n=10). The letters on top of the dietary (green) boxplots denote statistical differences between groups as determined by an ANOVA test followed by Tukey’s HSD post hoc comparison (p<0.05). Groups that have at least one similar letter are not significantly different from each other, while groups with different letters are significantly different. D) Boxplots showing the abundance of dietary proteins in fecal samples of conventional male (blue) and female (pink) mice. The asterisk indicates a significant difference between groups as determined with Welch’s T-test. The Benjamini-Hochberg procedure was used to correct for multiple hypothesis testing (p<0.05). E-J. PCA plots showing compositional differences between the fecal dietary proteomes of germ-free and conventional mice fed E) soy, F) casein, G) brown rice, H) yeast, I) pea, and J) egg white protein diets. Metaproteomic data used for A and C can be found in Supplemental Table 2.

Similar to the germ-free mice, the conventionally raised mice fed the 20% brown rice protein diet had the highest amount of dietary protein in the fecal samples, followed by the groups fed the 20% casein, 40% casein, and 20% egg white protein diet (Fig. 2C). We also analyzed differences in the abundance of dietary protein recovered in the fecal samples of male and female mice in both the germ-free and conventionally raised groups. Although there were no statistically significant differences in the abundance of dietary proteins recovered in the fecal samples of germ-free male and female mice in any of the diet groups, dietary protein abundance in the fecal samples of female mice fed the casein and brown rice protein diets trended higher than in the male mice (Fig. 2B). In the conventional mice, the amount of dietary protein recovered in the fecal samples of female mice fed the 20% casein protein diet was significantly higher than in the males (Fig. 2D). While it was not significantly different, the amount of dietary protein recovered in the fecal samples of the female mice fed the 20% brown rice protein diet trended higher than in the males.

To determine the impact of the gut microbiota on the digestion of dietary proteins from different sources we compared the composition of dietary proteins detected in the fecal samples of germ-free and conventionally raised mice. We found that the presence or absence of the gut microbiota significantly impacted the composition of the dietary proteins recovered in the fecal samples irrespective of dietary protein source in that there was a significant difference between the conventional and germ-free groups across all six 20% protein diets according to a PERMANOVA analysis (p<0.05) (Fig. 2 E-J). Additionally, we observed that the quantity of dietary protein did not significantly impact the composition of the dietary proteins recovered in the fecal samples of germ-free versus conventionally raised mice fed soy and casein protein diets because there were no significant differences between the 20% and 40% diet groups within these diets (Fig. 2 E and F). In summary, we found that depending on the source of dietary protein, varying amounts of dietary protein escape host digestion and are accessible to the gut microbiota.

### Proteins with functions potentially impacting the host or the microbiota escape host digestion and are consumed by the gut microbiota

To determine which constituent proteins from each of the six dietary protein sources escape host digestion, we compared the abundance of dietary proteins recovered in the fecal samples of germ-free mice to their abundance in diet samples and identified dietary proteins enriched in germ-free fecal samples. We further determined which dietary proteins escape host digestion and are depleted in the presence of the gut microbiota by comparing the abundance of dietary proteins recovered in the fecal samples of germ-free and conventionally raised mice. Most proteins showed no difference between germ-free and conventionally-raised mice or were more depleted in conventionally-raised mice, which likely indicates consumption by the microbiota. Interestingly, we also found some dietary proteins that were more abundant in the presence of a gut microbiota. The specific proteins that escape host digestion and are accessible to the gut microbiota in each of the six dietary protein sources are described below.

**<u>Soy:</u>** Of the 649 proteins detected in the 20% soy protein diet, on average 23 dietary proteins were detected in the fecal samples of the germ-free mice fed the 20% soy protein diet and 28 were detected in the fecal samples of the conventional mice. Some of the most abundant proteins in the soy protein diet, including the beta-conglycinin alpha and beta subunits, glycinin G4, and the 2S albumin protein, were significantly depleted in germ-free fecal samples compared to the diet, indicating that the host efficiently digests them (Fig. 3A). However, several dietary proteins, including the Kunitz trypsin inhibitor, seed linoleate 13S-lipoxygenase-1, and lipoxygenase, were significantly enriched in the fecal samples of germ-free mice compared to their abundance in the soy protein diet, indicating that they are not efficiently digested and absorbed by the host and are accessible to the gut microbiota.

**Figure 3.**
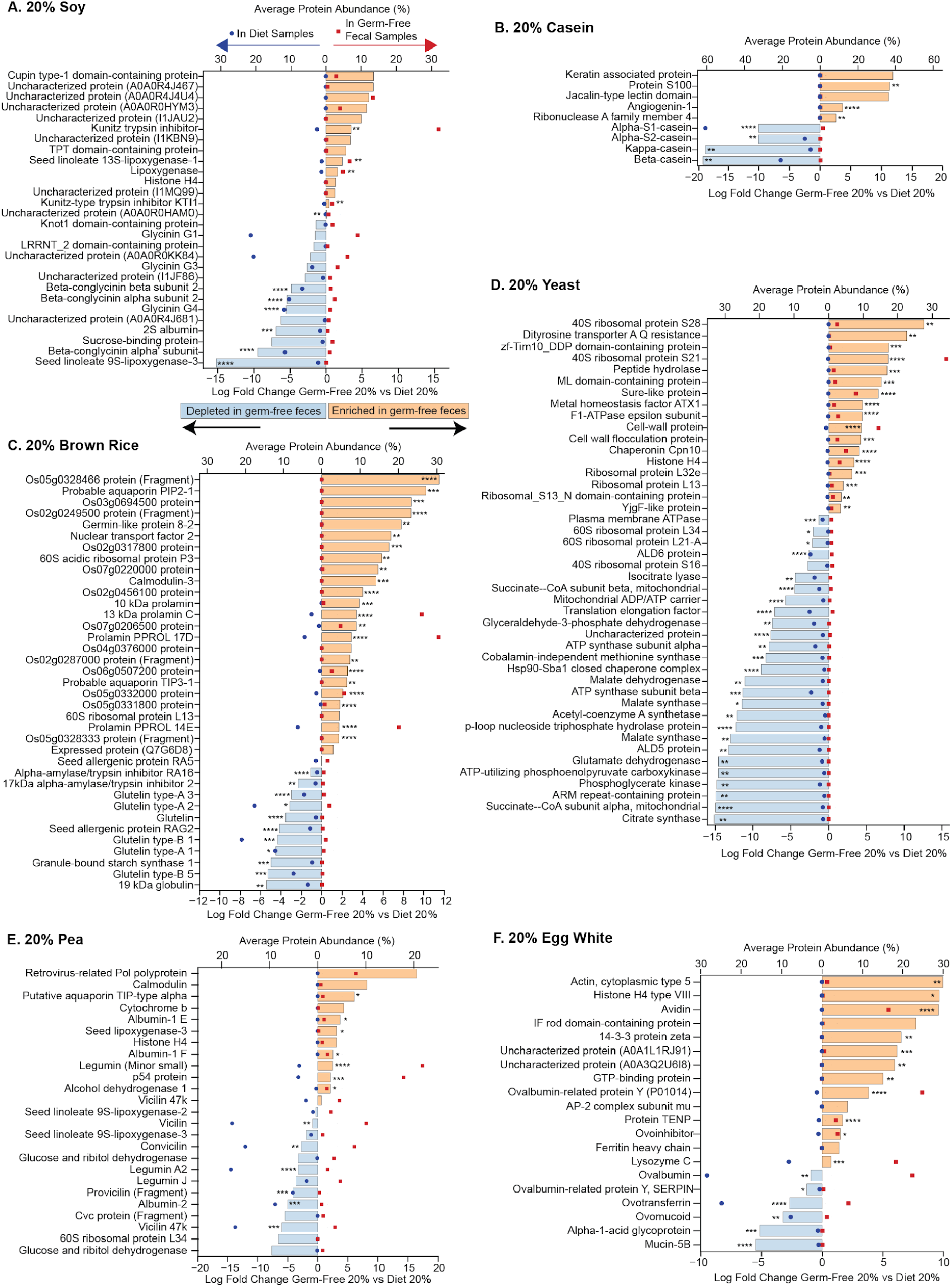
**Specific dietary proteins from different sources escape host digestion and are enriched in germ-free fecal samples at varying abundances**. A-F. Bar graphs showing the log_2_ fold change in abundance of dietary proteins in fecal samples of germ-free mice fed 20% soy, casein, brown rice, yeast, pea, and egg white protein diets as compared to the respective dietary samples. Dietary proteins enriched in germ-free fecal samples relative to the diet samples are depicted by the peach color bars and dietary proteins depleted in the germ-free fecal samples relative to the diet samples are depicted by the blue bars. The dynamic range of the LC-MS/MS approach used is 4 to 6 orders of magnitude and therefore displayed ratios of >10,000 fold are often due to the imputation of zeros in one condition with very small values to avoid infinite ratios and enable statistical testing. The overlaid scatter plot shows the average abundance of each protein in germ-free fecal samples (red squares) and dietary samples (dark blue circles). Average protein abundance refers to the mean abundance of the given protein across all replicates. The abundance of a given protein is relative to the abundance of all the other proteins in the dietary proteome. The asterisk indicates a statistically significant difference in the relative abundance of a dietary protein between the conventional and germ-free fecal samples as determined by Welch’s t-test. The Benjamini-Hochberg procedure was used to correct for multiple hypothesis testing (✱= p<0.05, ✱✱= p<0.01, ✱✱✱= p<0.001,✱✱✱✱= p<0.0001).Only proteins that were at least 1% abundant in either of the compared groups or had at least a 1 log fold difference are displayed. Significance was not used as a criterion for cutoff hence non-significant comparisons that meet the abundance and log-fold change cut-off are included. The complete data set and the correlating protein accession numbers can be found in Supplemental Table 3.

No dietary proteins were significantly depleted in the conventional fecal samples compared to the germ-free fecal samples (Fig. 4A); however, glycinin 1, glycinin 4, and sucrose-binding protein were enriched in conventional fecal samples as compared to germ-free fecal samples.

**Figure 4.**
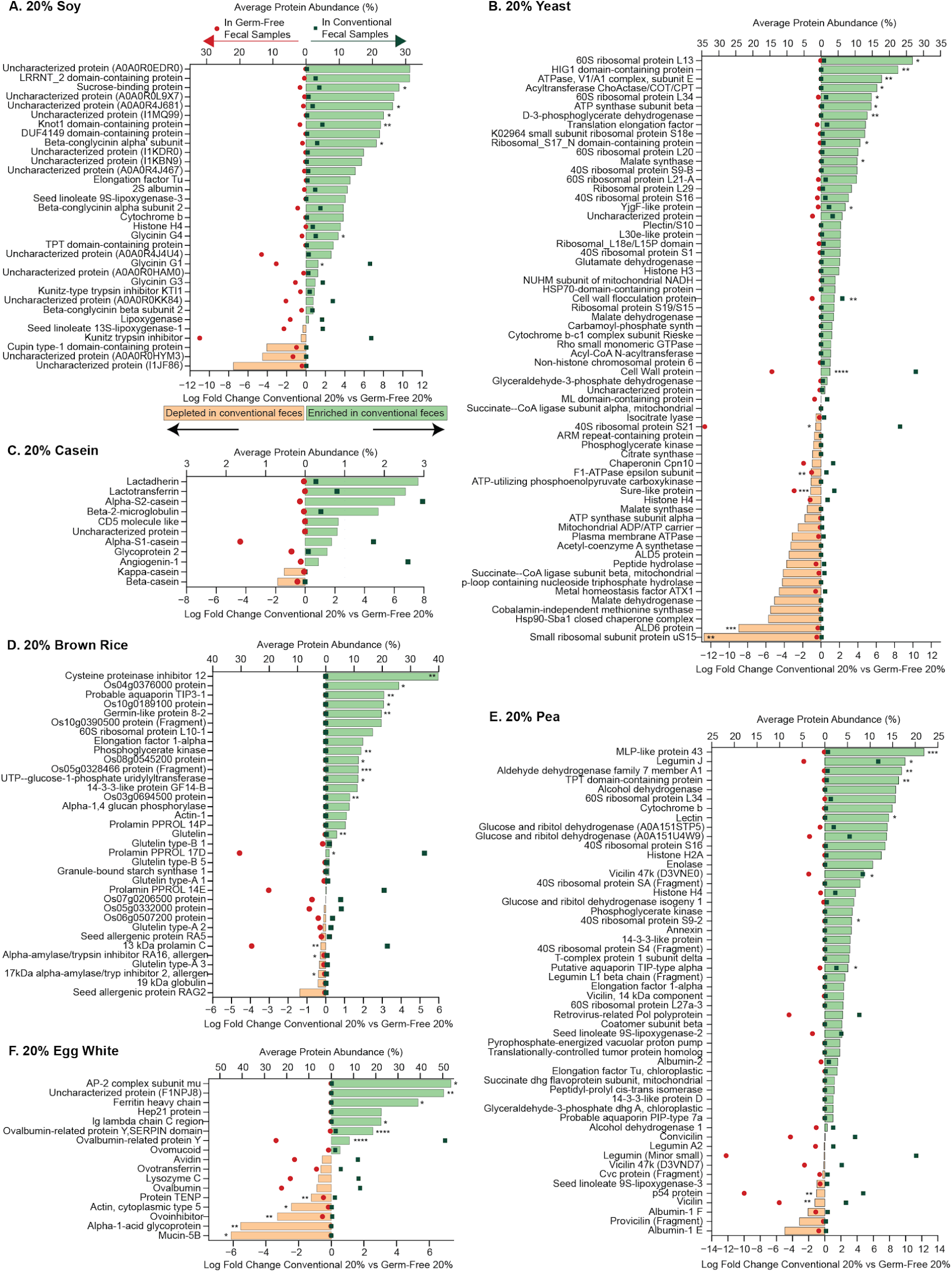
Specific dietary proteins that escape host digestion are consumed by the gut microbiota. A-F. Bar graphs showing the log_2_ fold change in abundance of dietary proteins in conventional versus germ-free mice fed 20% soy, casein, brown rice, yeast, pea, and egg white protein diets. Dietary proteins enriched in conventional fecal samples relative to the germ-free fecal samples are depicted by the green bars and dietary proteins depleted in the conventional fecal samples relative to the germ-free fecal samples are depicted by the peach bars. The dynamic range of the LC-MS/MS approach used is 4 to 6 orders of magnitude and therefore displayed ratios of >10,000 fold are often due to the imputation of zeros in one condition with very small values to avoid infinite ratios and enable statistical testing. The overlaid scatter plot shows the average abundance of each protein in conventional (dark green squares) and germ-free (red circles) fecal samples. Average protein abundance refers to the mean abundance of the given protein across all replicates. The abundance of a given protein is relative to the abundance of all the other proteins in the dietary proteome. The asterisk indicates a statistically significant difference in the abundance of dietary protein between the conventional and germ-free fecal samples as determined by Welch’s t-test. The Benjamini-Hochberg procedure was used to correct for multiple hypothesis testing (✱= p<0.05, ✱✱= p<0.01, ✱✱✱= p<0.001,✱✱✱✱= p<0.0001).Only proteins that were at least 1% abundant in either of the compared groups or had at least a 1 log fold difference are displayed. Significance was not used as a criterion for cutoff hence non-significant comparisons that meet the abundance and log-fold change cut-off are included. The complete data set and the correlating protein accession numbers can be found in Supplemental Table 3.

**<u>Casein</u>**: We detected 189 proteins (122 bovine, 67 *L. lactis*) in the 20% casein protein diet. Surprisingly, we detected more diet-derived proteins in fecal samples, 236 dietary proteins (20 bovine, 216 *L. lactis*) in the germ-fecal samples and 243 (19 bovine, 224 *L. lactis*) in the conventional fecal samples. In terms of relative abundance, while the *L. lactis* proteins constituted less than 1% of the purified protein diet, they comprised about 90% of the dietary proteins detected in the fecal samples of both germ-free and conventional mice (Supplemental Table 3). In terms of bovine proteins in the casein diet, the most abundant proteins in the diet, including alpha, beta, and kappa casein, were significantly depleted in the germ-free fecal samples indicating that they were efficiently digested by the host (Fig. 3B). Protein S100, angiogenin-1 and ribonuclease 4, were significantly enriched in the germ-free fecal samples indicating that they are not efficiently digested and absorbed by the host. When we analyzed fecal samples from germ-free mice fed a 40% casein protein diet, we detected the same proteins as in the 20% group but at a higher log fold change, which indicates that these proteins are digested and absorbed by the host regardless of their quantity.

Several of the *L. lactis* proteins in the casein diet that escaped host digestion and were enriched in the germ-free fecal samples were significantly depleted in the presence of the gut microbiota in the conventional fecal samples, including Beta-Ala-Xaa dipeptidase, triosephosphate isomerase, and DNA protection during starvation protein (Dps) (Supplementary Table 3). Bovine proteins like beta and kappa casein were also depleted in the presence of the gut microbiota, but the trends were not statistically significant (Fig. 4B). **<u>Brown Rice:</u>** On average, of the 205 proteins detected across all 20% brown rice protein diet samples, 74 proteins were detected in the germ-free fecal samples and 68 were detected in conventional fecal samples. In the germ-free fecal samples, these proteins included seed storage proteins such as 13 kDa prolamin C, prolamin 17D and prolamin 14E (Fig. 3C). Additionally, proteins of unknown function were enriched in the germ-free fecal samples relative to the diet. Three of these are characterized as seed storage proteins in UniProt (Q8GVK5, Q6ZIX4, Q5Z9M9), and one was characterized as a transmembrane protein (Q10ET9). Interestingly, several proteins were below the limit of detection in the diet but were detected in the fecal samples. These proteins included the aquaporin PIP2-1 protein, 60S acidic ribosomal protein, germin-like protein and 4 uncharacterized proteins. In contrast, other seed storage proteins like glutelin type proteins and 19 kDa globulin protein were significantly depleted in the germ-free fecal samples relative to diet, indicating that they are more efficiently digested and absorbed by the host than the prolamin type proteins.

Alpha-amylase/trypsin inhibitor RA16, 17kDa alpha-amylase/trypsin inhibitor 2, granule-bound starch synthase 1, and seed allergenic protein RAG2 were also efficiently digested by the host and were significantly depleted in the germ-free fecal samples compared to diet.

Compared to their abundance in the germ-free fecal samples, several seed storage proteins in the brown rice protein diet, including 17kDa alpha-amylase/trypsin inhibitor 2, alpha-amylase/trypsin inhibitor RA16, and 13 kDa prolamin C, were significantly depleted in the presence of the gut microbiota in the fecal samples of conventionally raised mice (Fig. 4C). Other dietary brown rice proteins like aquaporin TIP3-1, germin-like protein and a few uncharacterized proteins were significantly enriched in the conventional fecal samples compared to the germ-free fecal samples.

**<u>Yeast:</u>** Of the 1,476 dietary proteins identified in the yeast protein diet, we detected on average 77 proteins across all the germ-free fecal samples and 49 in the conventional fecal samples. Ribosomal proteins such as the 40S ribosomal protein S21, as well as cell wall proteins (CWP1 and FLO9), were significantly enriched in germ-free samples compared to their abundance in the diet, indicating that they were not efficiently digested and absorbed by the host (Fig. 3D).

In the conventional fecal samples, small ribosomal subunit protein uS15 (RPS13), ALD6 protein and SurE-like protein were significantly depleted in the presence of a gut microbiota compared to their abundance in the germ-free fecal samples (Fig. 4D).

Interestingly, the glycosylated cell wall proteins previously identified as resistant to host digestion were enriched in the conventional fecal samples.

**<u>Pea:</u>** Of the 511 proteins detected on average across the 20% pea protein diet samples, we detected 26 dietary proteins in the germ-free fecal samples and 34 in the conventional fecal samples. Seed storage proteins like albumin 1-E, albumin 1-F, and legumin were significantly enriched in the germ-free samples as compared to the diet, indicating that they are not efficiently digested by the host (Fig. 3E). Seed lipoxygenase was also significantly enriched in germ-free fecal samples, which was similar to the observed enrichment of seed lipoxygenase in the soy diet. Interestingly, the retrovirus-related Pol polyprotein and cytochrome b proteins were below the level of detection in the diet samples but were detected in the germ-free fecal samples. Vicillin, convicillin, provicillin, albumin 2, and legumin A2, also seed storage proteins in pea, were depleted in the germ-free fecal samples compared to the diet indicating that they are efficiently digested and absorbed by the host.

Vicillin and p54 protein were significantly depleted in fecal samples of conventionally raised mice compared to their abundance in the germ-free samples (Fig. 4E). Vicillin and p54 protein were also significantly enriched in germ-free fecal samples compared to diet, indicating that the host did not efficiently digest these two proteins in the pea protein diet and the gut microbiota subsequently had access to them. Other pea proteins like legumin J and lectin were significantly enriched in the fecal samples of conventionally raised mice compared to germ-free mice.

**<u>Egg White</u>**: Out of the 44 proteins detected on average in the 20% egg white protein diet, we detected 20 in the germ-free fecal samples and 18 in the conventional fecal samples. The top 4 most abundant proteins constituting 92% of the egg white diet were all detected in the fecal samples of germ-free mice, including ovalbumin, lysozyme C, ovomucoid, and ovotransferrin. Compared to their abundance in the diet, actin, avidin, ovalbumin-related protein Y, protein TENP, ovoinhibitor, and lysozyme C were significantly enriched in the germ-free fecal samples, indicating lower digestive efficiency for these specific proteins by the host (Fig. 3F).

In the fecal samples of conventionally raised mice, we found that heavily glycosylated proteins such as mucin-5B, alpha-1-acid glycoprotein, and ovoinhibitor were significantly depleted in the presence of the gut microbiota compared to their abundance in germ-free fecal samples (Fig. 4F). Additionally, antimicrobial proteins like protein TENP and lysozyme C were also depleted in the presence of a gut microbiota. Interestingly, Ovalbumin-related protein Y was enriched in the fecal samples of conventional mice compared to germ-free mice.

### Presence or absence of the gut microbiota does not impact dietary protein degradation by the host in the small intestine

To determine whether differences in dietary protein digestion observed between germ-free and conventional mice (Fig 2E-J, Fig 4) were due to microbial effects or differences in host digestive physiology, we compared dietary proteins in the small intestine (duodenum and ileum), large intestine (cecum and colon), and feces of germ-free and conventional mice fed the 20% soy and 20% casein diets. We found that the presence or absence of the gut microbiota did not impact dietary protein composition in the small intestine (duodenal and ileal contents) (Fig 5). Comparison of dietary proteins in the duodenal and ileal contents of conventional versus germ-free revealed no significant differences in both soy and casein groups, as determined by PERMANOVA analysis (p > 0.05). However, the presence or absence of the gut microbiota significantly impacted dietary protein content in the large intestine (cecal and colonic contents) and feces. Dietary proteins in the cecal contents of mice fed soy protein and in the colonic and fecal samples of mice fed soy or casein protein were significantly different between conventional and germ-free mice as determined by PERMANOVA analysis (p ≤ 0.05).

**Figure 5.**
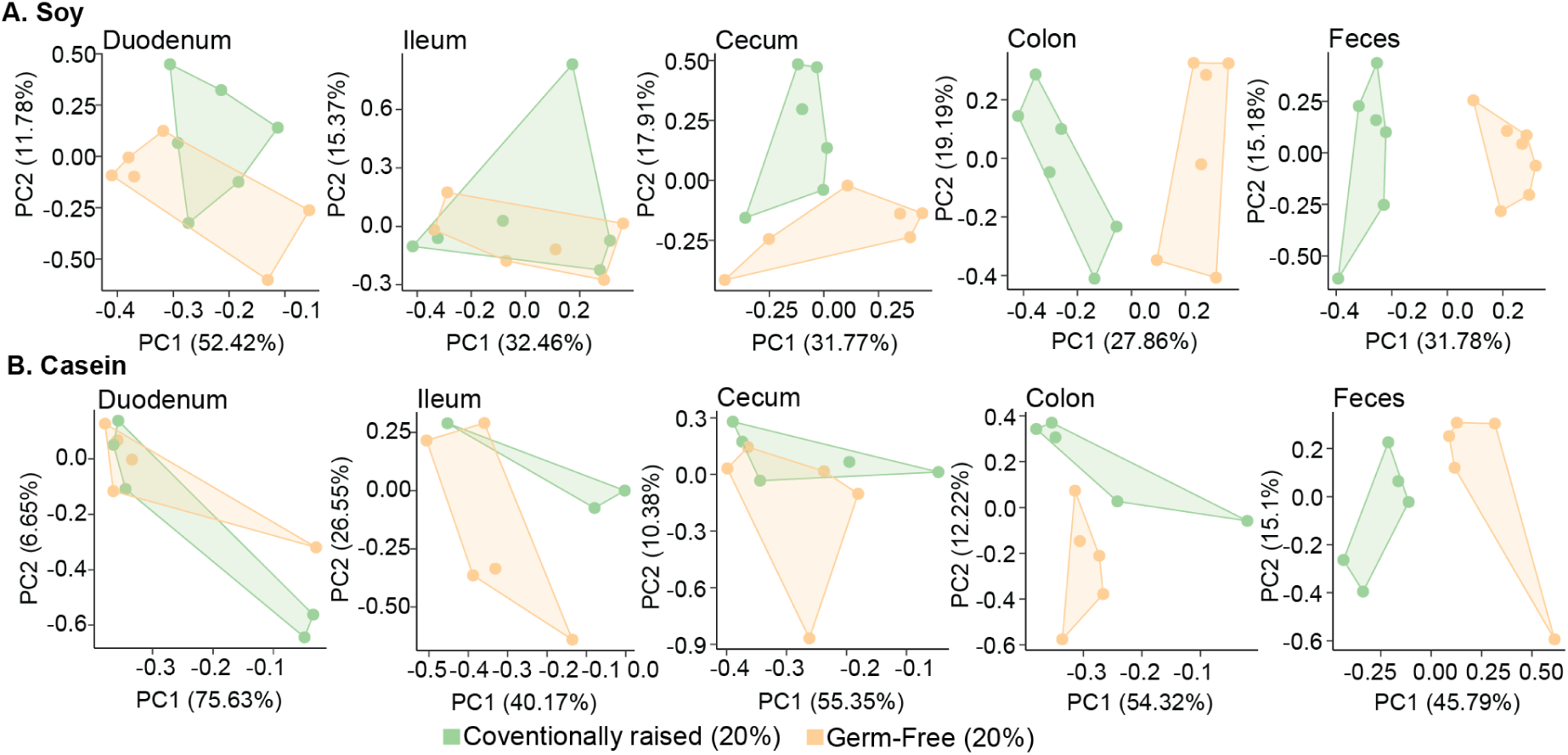
Presence or absence of gut microbiota does not impact dietary protein degradation in the small intestine but does in the large intestine. PCA plots showing compositional differences between the dietary proteins in duodenal, ileal, cecal, colonic and fecal contents of germ-free and conventional mice fed A) soy and B) casein. Underlying proteomic data can be found in Supplemental Table S6.

## Discussion

Diet impacts the host directly and indirectly via its interactions with the gut microbiota. Hence, it is crucial to study the dietary components impacting the composition and function of the gut microbiota, which can ultimately have consequences for host health. While these interactions are well studied in the case of dietary fiber and fat^29,32,33^, dietary protein has been investigated to a much lesser extent. In this study, we used a proteomics approach to profile the composition of dietary proteins from different sources and determined the fate of these dietary proteins in the host and their availability to the gut microbiota. We found that each of the dietary proteins from various plant and animal sources was unique in their composition and complexity. While the number of constituent proteins in animal sources like casein and egg was generally lower than in plant and microbial dietary proteins, the difference was not always significant (casein versus brown rice). Our results showed that the individual source of the dietary protein and its constituent proteins, rather than an overarching plant versus animal protein dichotomy, is more important in determining how the dietary protein is processed in the gut. For example, dietary proteins from brown rice were not efficiently digested, while other dietary proteins of plant origin like soy and pea were efficiently digested by the host.

Interestingly, we found that proteins from egg white, which has been shown to be highly digestible by current measures of digestibility, were detected in higher quantities in fecal samples of both germ-free and conventional mice as compared to the soy and pea proteins^34^. This indicates that egg white may not be as efficiently digested by the host as compared to these plant-based protein sources. Additionally, based on its protein digestibility-corrected amino acid score (PDCAAS), egg white is considered as high quality and efficiently digestible as casein and both have a score of 1.0 (highest possible score). PDCAAS is a protein quality metric that takes protein digestibility and amino acid composition and availability into account^13,16^. Our results, however, show that a significantly higher abundance of egg white proteins was recovered in the fecal samples of germ-free mice compared to casein. This may indicate that these two animal-sourced dietary proteins are digested differentially by the host and that egg white proteins may be less digestible and bioavailable than previously considered.

We further found that the *L. lactis* proteins in the casein protein diet were detected in high abundance in fecal samples despite the diets being sterilized using gamma irradiation. These results show that microbial proteins from *L. lactis* were not efficiently processed by the host. While the casein proteins in the diet were efficiently digested by the host, the L. lactis proteins escaped host digestion and were enriched in the fecal samples leading to their increased abundance in fecal samples compared to the diet samples. This observation raises important questions regarding how microbial proteins in our diets, especially fermented foods and alternative microbial-derived diets, are digested and processed by the host and the gut microbiota. Previous studies found that the gram-positive *L. lactis* present in casein was present in fecal samples of mice in the form of intact cells^29^. Based on these observations, it could be further hypothesized that other gram-positive microbes in fermented foods, such as *Streptococcus thermophilus* in yogurt, may also be detected in the form of intact cells in fecal samples after passage through the intestinal tract. On the other hand our results showed that eukaryotic microbial proteins from yeast were detected in much lower abundance in fecal samples compared to *L. lactis* proteins. This could be due to the fact that the yeast protein was processed and purified and likely no intact cells were present in the yeast protein diet allowing for more efficient digestion. These differences in how microbial proteins from different organisms are processed in the gut may have important implications for their nutritional relevance to the host and their impacts on the gut microbiota and host health.

Our results show that dietary proteins from all six sources were processed differently in the gut in the presence and absence of the gut microbiota. These differences indicate that the gut microbiota is involved in the degradation of these proteins. In a separate study we determined that the consumption of these undigested dietary proteins from different sources by the gut microbiota lead to strong effects on microbial composition and metabolism, specifically amino acid and glycan metabolism in mice fed the brown rice, yeast and egg white diets^11^. We identified hundreds of individual proteins in these sources that were resilient to host degradation and absorptive processes and were enriched in fecal samples. We also identified proteins that were differentially depleted only in the presence of the gut microbiota. This differential depletion only in the presence of the gut microbiota shows that these dietary proteins escaped host digestion, but were consumed by the gut microbiota.

Interestingly, we found several proteins, such as the beta-conglycinin alpha subunit in soy, lectin and legumin J in pea, ovalbumin in egg white, and cell wall protein in yeast, that were significantly enriched in the presence of gut microbiota. It is possible that these proteins were present in similar amounts in stool samples of germ-free and conventional mice but evaded detection by our mass spectrometry methods due to protein modification such as glycosylations. Microbial modification such as deglycosylation of dietary proteins in conventional mice may have made the proteins accessible for mass spectrometric detection in those samples. While this explanation would be partially due to detection limitations of the proteomic method it would still be indicative of a microbial influence on dietary protein digestion.

Specific dietary proteins that were not efficiently digested by the host included antinutritional factors such as the Kunitz trypsin inhibitor (KTI) in soy and lectin in pea protein. These proteins may impact the degradation and absorption of dietary components in the gut as trypsin inhibitors in plant proteins such as KTI have been shown to impact the proteolytic activity of digestive proteases like trypsin and chymotrypsin^35,36^. Furthermore, plant lectins can bind the epithelial cells in the digestive tract impacting the absorption of nutrients^37^. Additionally, several allergenic proteins including the vicillin seed storage protein in pea^38^, 17kDa alpha-amylase/trypsin inhibitor 2 and alpha-amylase/trypsin inhibitor RA16 in brown rice^39^, and ovalbumin, mucin-5B and ovoinhibitor in egg white^40,41^ were differentially depleted in the presence of the gut microbiota which may indicate that the gut microbiota plays a role in their degradation in the gut.

Several proteins in egg white with functions that could impact the host or the microbiota were not efficiently digested by the host, including avidin, ovalbumin, ovoinhibitor, and lysozyme C. Avidin strongly binds biotin and is used to induce biotin deficiency in dietary deficiency models. Avidin-induced deficiency of biotin has been shown to cause changes in gut microbiota composition and even induce IBD-like phenotypes^42^. This suggests that undigested avidin that reaches the large intestine can impact the gut microbiota and may play a role in gastrointestinal diseases. Based on our results it would be critical to study whether physiologically relevant amounts of avidin reach the large intestine in animals and humans after egg consumption and the level of activity of avidin within the gut environment. Ovalbumin has been shown to alleviate colitis symptoms in a DSS model of IBD^43^, and peptides produced from its hydrolysis have shown antihypertensive and antimicrobial properties^44^. Antimicrobial proteins that escape host digestion, such as lysozyme C in egg white, may alter gut microbial composition. Lysozyme C has been shown to be heat stable and retain its antimicrobial activity over a wide pH range^45^. Its presence in the fecal samples indicates that it can interact with the gut microbiota which can have implications for host health^46^. However, how different types of food processing techniques and digestive processes affect its activity in the gut and impact on the gut microbiota needs to be further investigated. In addition to these functionally relevant egg white proteins, we found that proteins with specific structural features such as glycosylations were differentially digested in the presence and absence of the gut microbiota. These included mucin 5B and alpha-1-acid glycoprotein in egg white and cell wall proteins in yeast. We previously showed that egg white and yeast protein diets impact the glycan metabolism of the gut microbiota and now our results suggest that these specific glycoproteins may be involved in affecting microbiota glycan metabolism^11,47^. In summary, our results show that dietary proteins from various sources with specific functional and structural features relevant to host physiology and microbial metabolism were differentially processed in the gut. However, further work is needed to determine whether these proteins remain functionally active in the gut as the detection of peptides here does not indicate whether the proteins are still in their native form or functionally active.

There are a few limitations to our study that should be further investigated in future studies. Our study only used purified dietary proteins in completely defined diets, which doesn’t account for the effects a complex food matrix would likely have on protein digestion and microbial interactions. Additionally, while we did not observe significant differences in the digestion of soy and casein between germ-free and conventional mice by the host in the small intestine, we did not examine how differences in digestive physiology between these groups might affect the breakdown of specific component proteins within these sources or other dietary sources beyond soy and casein^31,33^. Our study also does not investigate which specific microbial species consume which specific proteins or how these diet-microbiota interactions correlate with the host response. Future studies should investigate the functional activity of anti-nutritional factors and diet-derived antimicrobial proteins in the gut, how different ways of processing these dietary proteins before consumption affect the results, and the impact of specific dietary proteins identified as substrates of microbial metabolism on health and disease relevant members of the gut microbiota. Proteomics approaches can further be used to determine the digestibility of individual dietary proteins in food mixtures in real-life nutrition scenarios in humans.

Overall, our results provide an in-depth view of how the host and the microbiota impact the fate of different dietary protein sources in the gut and how individual component proteins within these sources are processed by the host. Additionally, we were able to determine how the gut microbiota impact the degradation of these proteins in the gut and identify specific dietary proteins that escape host digestion likely serving as substrates for gut microbial metabolism. These proteins, in some cases, have functions that could have impacts on health and physiology and should, therefore be further investigated in various human health conditions.

## Ethics statement

NC State’s Institutional Animal Care and Use Committee approved the animal care protocol (Protocol # 18-034-B for conventionally raised mice and # 18-165-B for germ-free mice).

## Data availability

Proteomics data, including raw mass-spectrometry files and databases used, were deposited to the ProteomeXchange Consortium via the PRIDE partner repository with the dataset identifier PXD041586.

## Supplemental materials

**Supplemental Table S1:** Composition of dietary proteins from Figure 1C. This table contains the accession numbers, abundance (percentage of proteome) and protein names of component proteins that make up the top 80% of each diet. “Other” in the Accession column is used to denote the abundance of all the lower abundance proteins that did not constitute the top 80% of the diets by protein abundance.

**Supplemental Table S2:** Table showing non-normalized abundance of all master proteins in all diet and fecal samples identified at 5% FDR. Naming convention: For fecal samples the first letter and number combination indicates the type of mouse (C=conventional, GF=germ-free) and cage number, followed by the number of the mouse tag in the cage, which is followed by the amount and type of protein diet. E.g C1_1_20SOY_Proteins.txt = ConventionalMouseCage1_MouseNumber1_20%SoyProteinDiet.”

**Supplemental Table S3:** Normalized abundance of dietary proteins for each protein source. Protein abundance within a sample was normalized using total sum scaling. Only proteins present in at least 75% of 1 group (mouse_type+diet) are included.

**Supplemental Table S4:** Composition of defined diets containing purified proteins from different sources.

**Supplemental Table S5:** Amino acid composition of defined diets containing purified dietary proteins from different sources.

**Supplemental Table S6:** Dietary proteins detected in the duodenal, ileal, cecal, colonic and fecal contents of germ-free and conventional mice fed soy and casein protein diets.

## Acknowledgments

We thank Dr. Heather Maughan for editing the manuscript and for her insightful suggestions. All LC-MS/MS measurements were made in the Molecular Education, Technology, and Research Innovation Center (METRIC) at North Carolina State University. The Gnotobiotic Core at the College of Veterinary Medicine, North Carolina State University is supported by the National Institutes of Health funded Center for Gastrointestinal Biology and Disease, NIH-NIDDK P30 DK034987. This work was supported by the National Institute Of General Medical Sciences of the National Institutes of Health under Award Number R35GM138362.

## Author contributions

**AA:** Data processing, data analysis, writing the manuscript, editing

**AB:** Conceptualization of the study, experimental design, data collection, editing

**JABR:** Data processing, editing

**TR:** Data collection

**CMT:** Experimental design, data collection, editing

**MK:** Funding, conceptualization of the study, experimental design, data collection, data processing, data analysis, writing, editing

